# A drug repurposing screen reveals dopamine signaling as a candidate therapeutic pathway for PIGA-CDG

**DOI:** 10.64898/2026.04.17.719256

**Authors:** Miriam C. Aziz, Jennie Wilson, Clement Y. Chow

## Abstract

PIGA-CDG is a congenital disorder of glycosylation caused by pathogenic partial loss-of-function variants in the *PIGA* gene. *PIGA* encodes an enzyme responsible for the catalytic transfer of N-acetylglucosamine to phosphatidylinositol during the first step of glycosylphosphatidylinositol anchor biosynthesis. Loss of this enzyme has a widespread phenotypic impact, but primarily results in neurological symptoms including seizures, intellectual disability, and developmental delay. Currently, treatments are limited and focus on symptom management. We developed an eye model of PIGA-CDG that has a reduced eye size. We screened a library of 98% 1,520 FDA/EMA-approved compounds to find drugs that improved the small eye phenotype. This screen revealed numerous drugs that improved eye size, including those that targeted dopamine signaling and cyclooxygenases. Using pharmacological and genetic approaches, we show that modulating dopamine signaling improves the eye size. Genetic inhibition of dopamine 2 receptor signaling and dopamine reuptake improve both the eye model and neurologically relevant PIGA-CDG phenotypes, including seizures and locomotor deficits. We also pharmacologically and genetically validate cyclooxygenase targeting drugs in the eye model. These findings reveal novel biology underlying PIGA-CDG and point towards candidate therapeutic approaches.

**AUTHOR SUMMARY:** PIGA-CDG is a rare neurodevelopmental disorder caused by pathogenic variants in the gene *PIGA*. Patients primarily display neurological symptoms, including seizures, developmental delay, and intellectual disability. Fewer than 100 patients have been identified, and treatment strategies are limited. In the context of rare diseases, *de novo* drug development is difficult due to the high cost, lengthy development times, and often too small of a patient population to conduct a clinical trial. Our lab leverages drug repurposing screening to circumvent many of the hurdles associated with *de novo* drug development. Here, we develop and screen FDA- or EMA-approved compounds on a *Drosophila* model of PIGA-CDG, uncovering novel biology underlying *PIGA*-associated pathophysiology. We use pharmacological and genetic tools to demonstrate that modifying dopamine signaling and abundance, as well as cyclooxygenase-mediated pathways, contribute to *PIGA* associated phenotypes. This work highlights promising therapeutic targets for PIGA-CDG.

## INTRODUCTION

PIGA-CDG is an ultra-rare X-linked recessive neurodevelopmental disorder caused by partial loss-of-function variants in the *Phosphatidylinositol glycan biosynthesis class A* (*PIGA*) gene [1–4]. Fewer than 100 patients have been identified with PIGA-CDG. Of these patients, approximately 40 partial loss-of-function variants have been reported [3]. Like other congenital disorders of glycosylation (CDGs), PIGA-CDG patients typically present with seizures, intellectual disability, developmental delay, hypotonia, dysmorphic facial features, and lower life expectancy [5]. Other organ systems can be affected, leading to skin abnormalities, gastrointestinal issues, respiratory problems, and other complications. There is no disease-specific treatment for PIGA-CDG and only symptom management is available. With a small patient population and limited funding, developing novel therapies and conducting clinical trials for rare diseases like PIGA-CDG is a significant challenge.

*PIGA* encodes a critical enzyme required for the first step of Glycosylphosphatidylinositol (GPI) anchor biosynthesis, a glycosylation process that is highly conserved from yeast to mammals [6–8]. PIGA is the essential catalytic subunit of the GPI-N-acetylglucosaminyl transferase complex. During the first step of GPI-anchor biosynthesis, PIGA transfers an N-acetylglucosamine (GlcNAc) from 5′-diphospho N-acetylglucosamine (UDP-GlcNAc) to phosphatidylinositol (PI) on the cytoplasmic side of the endoplasmic reticulum (ER) membrane, generating GlcNAc-PI, the first intermediate of GPI-anchor biosynthesis [9, 10]. Additional sugar and lipid moieties are sequentially added to GlcNAc-PI, generating a mature GPI anchor that is then linked to various proteins being synthesized in the ER, and ultimately sent to the cell surface [11]. Over 150 different GPI-anchored proteins (GPI-APs) are synthesized in the body, each contributing to diverse cellular functions, including signaling, adhesion, immunity, and more [5]. Due to the wide range of GPI-AP functions, it is challenging to determine how the loss of PIGA and subsequent depletion of GPI-APs contribute to the complex phenotypes observed in individuals with PIGA-CDG.

It is challenging to develop novel therapeutics for rare diseases, such as PIGA-CDG, due to a lack of funding, lengthy development times, and the limited number of patients required to conduct proper clinical trials. Current treatment strategies for PIGA-CDG are limited and primarily treat individual symptoms. Moreover, it is unclear why PIGA-CDG patients exhibit such severe clinical presentations, as the pathophysiology of PIGA-CDG is not well understood. In this study, we utilize a drug repurposing screen to rapidly identify compounds that are U.S. Food and Drug Administration (FDA)- or European Medicines Agency (EMA)-approved, enabling their rapid translation from bench to bedside. Additionally, by knowing the targets of these drugs, we can begin to understand the underlying pathophysiology behind disorders like PIGA-CDG.

Our lab has successfully utilized drug repurposing to identify candidate therapies and molecular targets for multiple *Drosophila* models of rare diseases, including DPAGT1-CDG and NGLY1-deficiency [12, 13]. *Drosophila* are a powerful tool for studying human disease, as they are low-cost, have short generation times, and share ∼75% disease-causing genes with humans [14]. Their short life cycle makes them especially useful for high-throughput screening. There are currently very few therapies that address the broad spectrum of symptoms seen in PIGA-CDG patients. One way to accelerate therapeutic discovery is to screen libraries of drugs that are FDA/EMA-approved, allowing for rapid translation from fly models to patients while bypassing the lengthy timelines and high costs associated with *de novo* drug development.

In this study, we conducted a drug repurposing screen using a *Drosophila* PIGA-CDG model to identify novel therapeutics and disease-relevant pathways. Of the 1,520 small molecules screened, we identified 89 compounds that suppressed the PIGA-CDG model phenotype, including those targeting dopaminergic, cyclooxygenase (COX), adrenergic, and steroid pathways, among others. Here, we demonstrate that both pharmacological and genetic inhibition of COX improves our PIGA-CDG eye model, supporting COX inhibition as a potential therapeutic strategy. We use pharmacological and genetic manipulation to show that dopamine signaling and dopamine abundance modifies developmental and behavioral PIGA-CDG associated phenotypes. Together, these results provide a strong foundation for defining the mechanisms underlying PIGA-CDG phenotypes and for advancing our understanding of novel therapeutic targets.

## RESULTS

### A drug repurposing screen reveals suppressors of a *Drosophila* PIGA-CDG eye model

To perform a drug repurposing screen, we developed an RNA interference (RNAi) based loss-of-function model of PIGA-CDG in *Drosophila* using the GAL4/UAS system. *PIGA* is knocked down only in eye tissue using the *eya-GAL4* driver [15]. *PIGA* knockdown in the eye results in a smaller eye with a glassy appearance (Fig 1A). Previously, we demonstrated that this same RNAi results in a 60% reduction in *PIGA* expression [16]. Because ubiquitous *PIGA* knockdown is lethal, a drug screen can only be performed in a tissue-specific knockdown model [16]. The *Drosophila* eye is a complex organ primarily composed of neurons, with mechanisms that overlap those in the brain, making it a suitable phenotypic model for PIGA-CDG. The small eye size also offers a valuable quantitative phenotype for high-throughput screening (Fig1A). Our lab uses this eye phenotype and pharmacological or genetic perturbations to investigate pathways and small molecules that can modify the severity of different disease models [12, 17–20].

**Fig 1:**
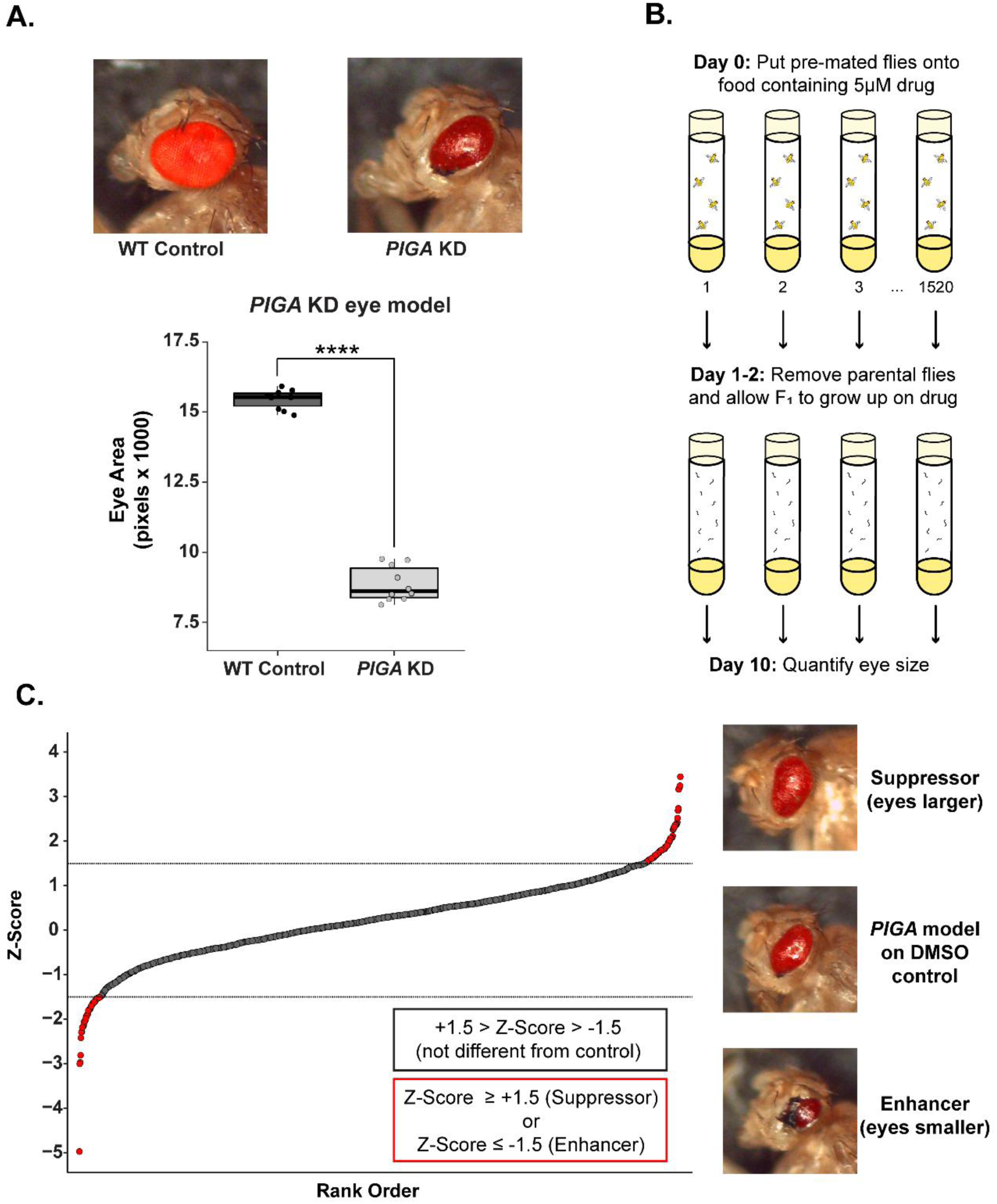
Drug repurposing screen pipeline and results. **(A)** Development of *PIGA* eye model. WT Control on top left and *eya*-GAL4 *PIGA* knockdown (KD) model on top right. Eye sizes are quantified in plot below. Statistical significance was determined using an unpaired two-tailed t-test. **** p<0.0001. **(B)** Schematic of drug repurposing pipeline. Flies were raised on 5 µM drug or 0.05% DMSO in standard food. Eye size of adults was quantified at two days post eclosion. (C) Z-Score plot of all drugs tested. Each point represents the Z-Score from each of the 1,520 drugs in the Prestwick Chemical library compared to DMSO control. We identified 89 suppressors with a Z-Score ≥ 1.5 and 56 enhancers with a Z-Score ≤ -1.5 (dotted lines). Representative images of the *PIGA* KD eye model on DMSO control and change in eye size when on suppressor or enhancer compounds.

Using the *PIGA* eye model, we performed a drug repurposing screen using the Prestwick Chemical Library (Fig 1B), a collection of 1,520 drugs, 98% of which are FDA/EMA-approved. We raised the *PIGA* eye model on 5 µM of drug and collected flies upon eclosion. We imaged and quantified the eye size of up to 5 flies on a drug and calculated a Z-Score by comparing to the average eye size to the *PIGA* eye model raised on DMSO (Fig 1B). We identified 89 compounds that increased eye size (suppressors, Z ≥ 1.5), 56 compounds that decreased eye size (enhancers, Z ≤ -1.5), and 41 compounds that resulted in complete lethality of the model (Fig 1C, S1 Table).

We prioritized validating drug classes that were recurrent in our suppressor set. Our primary screen revealed suppressors that fell into distinct categories, including six drugs that targeted dopamine signaling, six targeting cyclooxygenase (COX) enzymes, eight acting on adrenergic receptors, and six classified as steroids. We were most interested in therapeutics that were easy to translate to the clinic (i.e., available over-the-counter), safe for pediatric use, and targeted pathways relevant to patients’ symptoms (i.e., neurological).

### COX inhibition improves PIGA-CDG eye model

Cyclooxygenases (COX) are enzymes responsible for synthesizing prostaglandins, a group of lipids that function in inflammation, vasoconstriction/dilation, and platelet aggregation [21]. Non-steroidal anti-inflammatory drugs (NSAIDs) inhibit COX-1 and/or COX-2 enzymes (encoded by *PTGS1* and *PTGS2* in humans), thus inhibiting prostaglandin synthesis and providing anti-inflammatory relief. Our primary screen revealed six COX inhibitors that suppressed the eye phenotype: Naproxen, Fenclofenac, Aceclofenac, Sulindac, Etoricoxib, and Piroxicam. COX inhibitors are commonly used, and some do not require a prescription, making them easily accessible to patients. Naproxen, available over the counter, significantly increased eye size at all doses tested (9.3%, 15.7%, and 11% increases at 1 µM, 5 µM, and 25 µM respectively) (Fig 2A). Fenclofenac increased eye size at 1 µM (15.8% increase) and 5 µM (13.5% increase) doses (Fig 2B). Aceclofenac increased eye size at 25 µM (13.5% increase) (Fig 2C).

**Fig 2:**
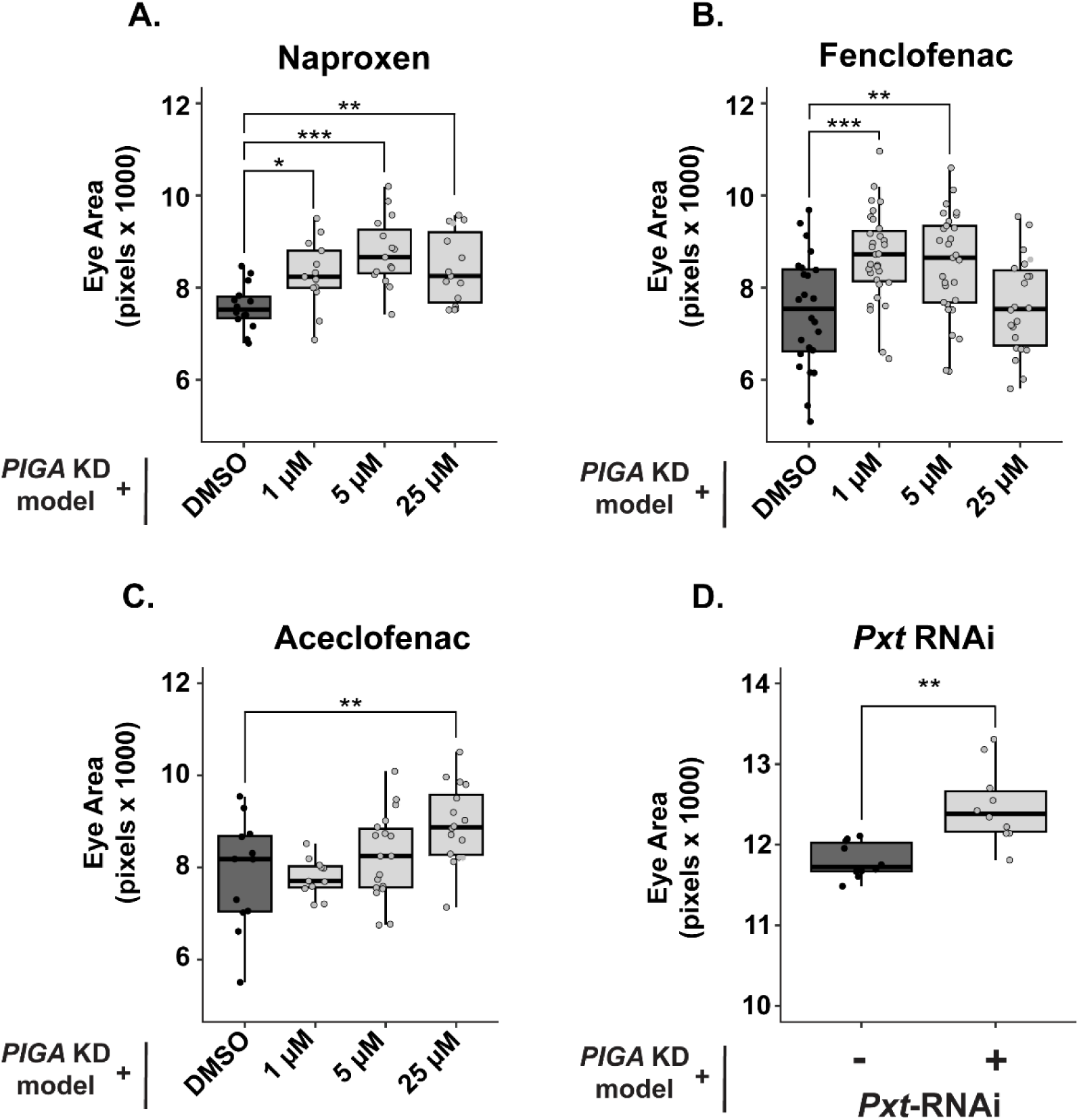
Cyclooxygenase inhibition improves the *PIGA* eye model. **(A-C)** Secondary validation of COX inhibitors identified in the primary screen. Naproxen increases eye size at 1 µM, 5 µM, and 25 µM by 9.3%, 15.7%, and 11%, respectively. Fenclofenac increases eye size at 1 µM and 5 µM by 15.8% and 13.5%, respectively. Aceclofenac increases eye size at 25 µM by 13.5%. **(D)** RNAi against COX-like gene *Pxt* (BDSC 28934) partially rescues eye size of *PIGA* eye model, increasing eye size by 7.2%. Statistical significance was determined using one-way ANOVA with Dunnett’s multiple comparisons test for drug treatments and unpaired two-tailed t-tests for genetic experiments. * p<0.05, ** p<0.01, *** p<0.001.

The *Drosophila* ortholog of human *PTGS1* and *PTGS2* is *Peroxinectin-like (Pxt)* [22]. Pxt is responsible for prostaglandin synthesis in the fly. We knocked down *Pxt* in the *PIGA* eye model and found that this also increased eye size (7.2% increase), similar to the effect of treatment with NSAIDs (Fig 2D, S1 Fig). These data suggest that inhibition of prostaglandin synthesis may be a therapeutic avenue for PIGA-CDG patients.

### Inhibition of Dopamine 2 Receptor signaling improves the PIGA-CDG eye model

We prioritized validating dopamine signaling targeting drugs because PIGA-CDG symptoms are primarily neurological. Our primary screen identified six dopamine signaling-targeting drugs: Quetiapine Hemifumarate, Octoclothepin Maleate Salt, Mesoridazine Besylate, Bromopride, Ziprasidone, and Alizapride hydrochloride. All these drugs are described as dopamine D2 receptor antagonists. The structure of the dopamine targeting drugs fell into two categories: tricyclics or non-tricyclics. Interestingly, tricyclic drugs partially rescued eye size in validation experiments, whereas the non-tricyclics failed to do so. This structure may be important for the partial rescue seen in the *PIGA* eye model. The tricyclic drugs that increased eye size in validation experiments include Quetiapine Hemifumarate, Octoclothepin Maleate Salt, and Mesoridazine Besylate. Quetiapine Hemifumarate, an atypical antipsychotic with affinity to D2R and serotonin receptors (5-HT2), increased eye size in male flies at 25 µM (12.9% increase), with small increases at 1 µM and 5 µM (7% and 6.8%, respectively) (Fig 3B). Octoclothepin Maleate Salt, an antipsychotic with affinity for D2R and 5HT-2A receptors, significantly increased eye size at 50 µM (9.4% increase) (Fig 3C). Mesoridazine Besylate, a tricyclic first-generation antipsychotic that blocks D2R and D4R signaling, increased male eye size by 8.4% at 25 µM. This consistent increase in eye size with D2R antagonist treatment supports the idea that Dop2R inhibition may offer broader therapeutic potential than captured in our initial assays.

**Fig 3:**
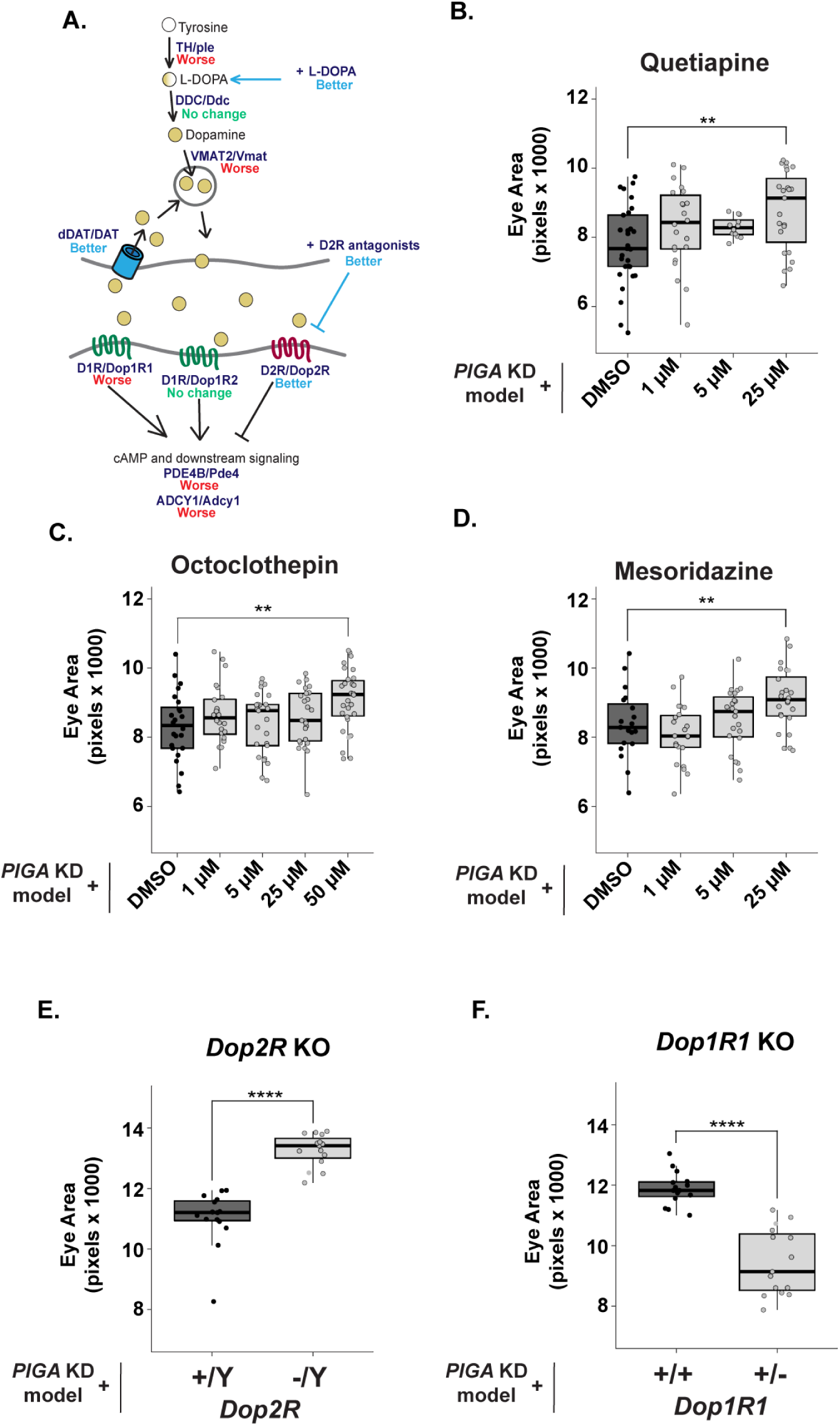
Pharmacological and genetic inhibition of Dopamine 2 receptor improves *PIGA* eye model. **(A)** Schematic of dopamine receptor signaling in humans/Drosophila. Effect of dopamine signaling pathway mutants on the *PIGA* eye model denoted in blue (better), red (worse), or green (no change). **(B-D)** Secondary validation of Dopamine 2 receptor antagonists identified in the primary screen. Quetiapine Hemifumarate increases eye size at 25 µM by 12.9%. Octoclothepin Maleate Salt increases eye size at 50 µM by 8.4%. Mesoridazine Besylate increases eye size at 25 µM by 9.4%. **(E)** Hemizygous loss of *Dop2R* (BDSC 84720) partially rescues the *PIGA* eye model, increasing eye size by 18.3%. **(F)** Heterozygous loss of *Dop1R1* (BDSC 92640) worsens the *PIGA* eye model, reducing eye size by 16.9%. Statistical significance was determined using one-way ANOVA with Dunnett’s multiple comparisons test for drug treatments and unpaired two-tailed t-tests for genetic experiments. ** p<0.01, *** p<0.001, **** p<0.0001.

The dopamine signaling targeting drugs that validated all antagonize the D2R receptor. In *Drosophila*, *Dop2R* is orthologous to mammalian *D2R, D3R,* and *D4R,* and inhibits cAMP signaling when activated [23, 24]. We next asked whether genetic inhibition of Dop2R would phenocopy the partial rescue observed with pharmacological inhibition of Dop2R. We introduced a hemizygous loss-of-function allele [25] (*Dop2R* is X-linked in *Drosophila*) into the *PIGA* eye model. Genetic loss of *Dop2R* significantly increased *PIGA* eye size with a much larger effect than observed with drug treatment (18.3% increase) (Fig 3D). These data support pharmacological data and highlight D2 receptor inhibition as a potential therapeutic strategy for PIGA-CDG.

In both *Drosophila* and humans, dopamine receptor signaling via dopamine 1 and dopamine 2 receptors has opposing downstream effects [23]. When D2R is activated, adenylate cyclase and cyclic adenosine monophosphate (cAMP) signaling are inhibited (Fig 3A). Activation of D1R stimulates downstream cAMP signaling (Fig 3A). In *Drosophila, Dop1R1* and *Dop1R2* are orthologous to *D1R* and *D5R* [23, 24]. We introduced a heterozygous null allele of *Dop1R1* [26], and as expected, this enhanced the phenotype and decreased eye size in the *PIGA* eye model (17.8% decrease) (Fig 3E). These results are consistent with the biology of dopamine receptor signaling and opposing effects on cAMP signaling. We did not see a change in the eye phenotype when introducing a heterozygous null allele of *Dop1R2* into the *PIGA* eye model, which could suggest that Dop1R1 signaling compensates for loss of Dop1R2.

### Increasing dopamine levels improve the PIGA-CDG eye model

Dopamine signaling begins with the synthesis of the dopamine precursor, L-DOPA. L-DOPA is produced through the hydroxylation of L-tyrosine by the enzyme *ple* (*tyrosine hydroxylase (TH)* in mammals) [23]. This reaction represents the first and rate-limiting step in dopamine synthesis. Levodopa (L-DOPA) is FDA-approved and commonly used to treat Parkinson’s disease. To test whether dopamine synthesis might impact PIGA-CDG, we raised the *PIGA* eye model on L-DOPA to stimulate dopamine synthesis. Strikingly, L-DOPA treatment partially rescued *PIGA* eye size (19.6% increase at 25 µM and 14% increase at 50 µM) (Fig 4A). Based on this, we hypothesized that inhibiting dopamine synthesis would have the opposite effect, enhancing the *PIGA* eye phenotype.

**Fig 4:**
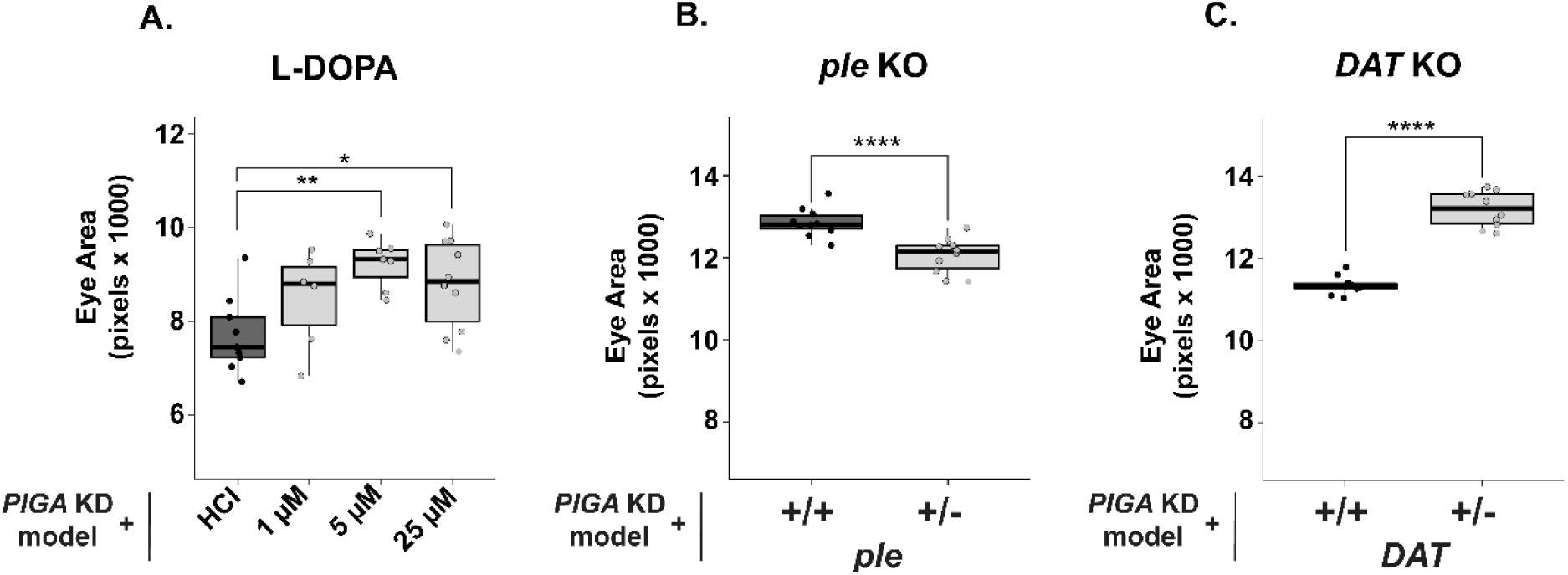
Increasing dopamine levels improves *PIGA* eye model. **(A)** Supplementing food with L-DOPA improves the *PIGA* eye model, increasing eye size by 19.6% at 25 µM and 14% at 50 µM. **(B)** Introducing a heterozygous null allele of *ple* (DGGR 106-955), which encodes the enzyme responsible for converting tyrosine to L-DOPA, decreased eye size by 6.3%. **(C)** A heterozygous null allele of *DAT* (BDSC 30867), a gene that encodes a dopamine recycling enzyme, increases eye size by 16.8%. Statistical significance was determined using one-way ANOVA with Dunnett’s multiple comparisons test for drug treatments and unpaired two-tailed t-tests for genetic experiments. * p<0.05, ** p<0.01, *** p<0.001, **** p<0.0001.

To test this, we introduced loss-of-function alleles in dopamine synthesis genes into the *PIGA* eye model. We introduced a heterozygous *ple* null allele [27] into the *PIGA* eye model. As expected, loss of *ple* enhanced the phenotype and made the eye size smaller (6.3% decrease) (Fig 4B). Interestingly, introducing a heterozygous null allele of the subsequent dopamine synthesis enzyme, *Ddc*, did not change eye size (data not shown). *ple* is likely more sensitive to loss because converting tyrosine to L-DOPA is the rate-limiting step in dopamine synthesis [28]. Strikingly, the mammalian ortholog of *ple, tyrosine hydroxylase (TH),* is the second most upregulated gene in a mouse model of PIGA-CDG [29]. *TH* may be upregulated to compensate for deficiencies in L-DOPA synthesis and downstream dopamine signaling.

We also found that the genetic inhibition of *Vmat* enhanced the eye phenotype to a greater extent than when we inhibited *ple* (22.4% decrease) (S2A Fig). Vmat transports dopamine and other monoamine neurotransmitters from the neuronal cytoplasm into synaptic vesicles. Accordingly, loss of Vmat disrupts the vesicular packaging of dopamine as well as serotonin, histamine, and octopamine, limiting dopamine-specific interpretation of this manipulation. Despite the broad monoaminergic effects of Vmat inhibition, the consistent enhancement of the phenotype following reduced dopamine synthesis underscores the sensitivity of this model to dopamine availability.

We next asked whether altering dopamine clearance could similarly impact the phenotype. We introduced a heterozygous null allele in the *dopamine active transport gene*, *DAT* [30], into the *PIGA* knockdown model and found that loss of DAT increased the eye size (16.3% increase) (Fig 4C). The inhibition of DAT may partially rescue the phenotype by allowing dopamine to remain in the synaptic cleft longer to bind to its receptors. Taken together, these findings provide compelling evidence that dopamine availability, either by increasing dopamine synthesis or reducing synaptic clearance, is a key modulator of disease severity in the *PIGA* eye model, highlighting dopamine synthesis as a potential therapeutic target for PIGA-CDG.

### Inhibition of cAMP phosphodiesterase and adenylate cyclase activity enhances the PIGA-CDG eye model

Dopamine receptors are G-protein-coupled receptors that differentially regulate intracellular cAMP [24]. Upon stimulation, Dop1R1 and Dop1R2 receptors are coupled to G_s_, which stimulates cAMP production by a Ca^2+^/Calmodulin stimulated adenylate cyclase (AC). Activation of Dop2R results in coupling to G_αi/o_, which inhibits adenylate cyclase and prevents cAMP synthesis. Upon cAMP increase, protein kinase A (PKA) is activated and phosphorylates downstream substrates such as cAMP response element-binding protein (CREB) [31]. Phosphodiesterases then degrade cAMP to maintain homeostasis. Core components of cAMP signaling such as the adenylyl cyclase *Adenylate cyclase 1 (Adcy1)* and the cAMP phosphodiesterase *Phosphodiesterase 4 (Pde4)* are essential regulators of cAMP levels in fly neurons and are implicated in associative behaviors linked to dopaminergic modulation [24, 31, 32].

Because loss of Dop2R partially rescued the *PIGA* eye model and Dop2R activation inhibits cAMP signaling, we hypothesized that loss of Dop2R would increase cAMP production, which could be the mechanism underlying the partial rescue observed in the eye model. To investigate how modifying downstream cAMP signaling impacts the *PIGA* eye model, we introduced the *Pde4* hypomorphic allele, which increases cAMP levels compared to wild type [33]. Hemizygous loss of *Pde4* (X-linked) in the *PIGA* eye model exacerbated the eye phenotype, resulting in a smaller eye (22.1% decrease) (S2B Fig). We also introduced a hemizygous null allele of *Adcy1* (X-linked) [34] into the *PIGA* eye model, which also resulted in a decrease in *PIGA* eye size (30.6% decrease) (S2C Fig). These data suggest that the role of cAMP in *PIGA* pathophysiology is likely more complex when considering the requirement of cAMP signaling across multiple cell types and neurotransmitter pathways. Pde4 also acts in the serotonin pathway, and loss of Pde4 in multiple neurotransmitter circuits may be detrimental to *PIGA* pathophysiology.

### Inhibition of Dopamine 2 Receptor signaling significantly improves *PIGA*-associated neurological dysfunction

Movement and muscle dysfunction are common amongst PIGA-CDG patients, phenotypes we can recapitulate in the lab. Pan-neuronal knockdown of *PIGA (elav*-GAL4) results in severe locomotor and neuromuscular defects, as well as a reduced lifespan [16]. Within 5 days following eclosion, most pan-neuronal *PIGA* knockdown flies develop a neuromuscular defect, causing an erect wing phenotype. Nearly all pan-neuronal *PIGA* knockdown flies have a severe locomotor deficit as measured by a negative geotaxis assay (S1 Video). Focusing on genes whose loss rescued the eye phenotype, we introduced the *Dop2R* null allele into the pan-neuronal model to determine whether loss of *Dop2R* could rescue multiple *PIGA*-associated neurological phenotypes. The locomotor defect is so severe in this model that flies struggle to even get off the bottom of the vial. After being tapped down, we measured the number of flies that left the bottom after 30 seconds of climbing. Only ∼10% of *PIGA* knockdown flies can climb (S1 Video). Due to the severe impairment in male flies, we only tested female flies for this model. When *Dop2R* is lost in the *PIGA* knockdown model (heterozygous loss, females tested), ∼25% of flies leave the bottom (Fig 5A, S2 Video), suggesting that loss of *Dop2R* is beneficial for PIGA-related movement phenotypes. Loss of *Dop1R1* did not significantly worsen the *PIGA* climbing phenotype, likely because flies that are already so severely impaired have limited capacity for further decline. We also investigated whether heterozygous loss of *DAT* would improve the *PIGA* pan-neuronal knockdown climbing model, however loss of DAT had no effect on the model (S3 Fig).

**Fig 5:**
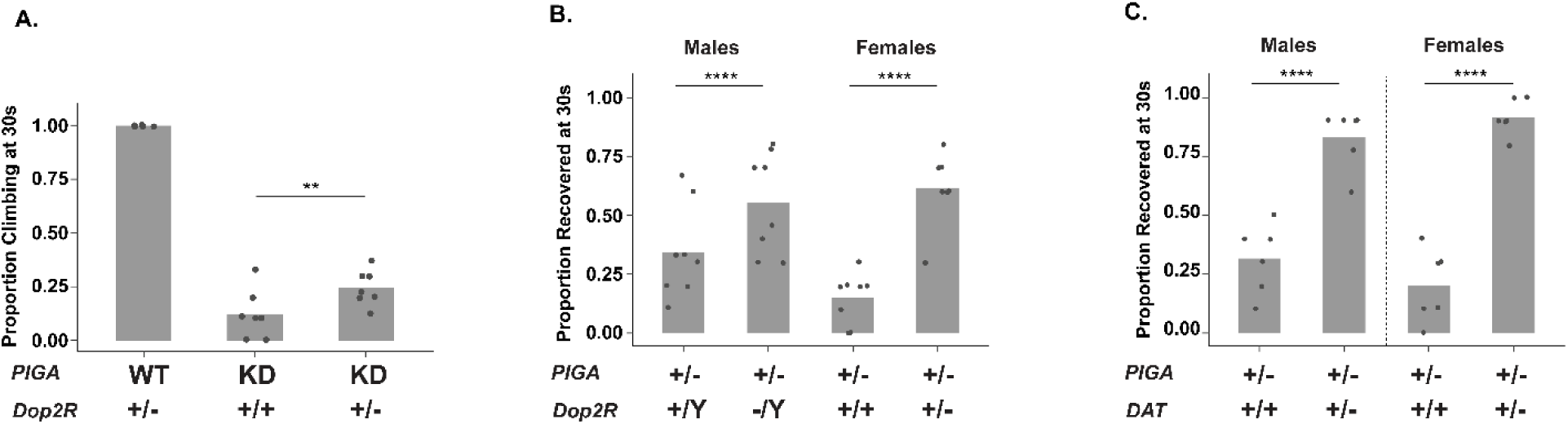
*Dop2R* loss improves *PIGA*-associated behavioral phenotypes. **(A)** Heterozygous loss of *Dop2R* improves the severe climbing impairment in pan-neuronal *PIGA* knockdown female flies. Only females were tested, as males were too severely impaired to be properly assessed. **(B)** Loss of *Dop2R* in PIGA null heterozygous flies increases the proportion of flies recovering from seizure activity after mechanical stimulation in a bang sensitivity assay. **(C)** Loss of *DAT* in PIGA null heterozygous flies increases the proportion of flies recovering from seizure activity after mechanical stimulation in a bang sensitivity assay. Each point represents an individual replicate of 8-11 flies. Statistical significance was assessed using a Chi-squared (χ²) test.

All PIGA-CDG patients have epilepsy, and most are refractory to current anti-seizure medications, demonstrating a need for identifying novel therapeutic approaches. We previously developed a PIGA-CDG model that has severe seizures caused by heterozygous loss of *PIGA* [16]. Homozygous loss of *PIGA* is lethal, but heterozygous null *PIGA* flies survive into adulthood, albeit with severe seizure phenotypes. Recovery following mechanical stimulation in the bang sensitivity assay is exceptionally prolonged in *PIGA* heterozygous null flies, with both males and females often taking more than three minutes to regain posture and leave the bottom of the vial. Using the same Dop2R antagonists that partially rescued eye size, we tested whether pharmacological Dop2R inhibition would improve seizure phenotypes. Pharmacological Dop2R inhibition had no effect on seizure severity (data not shown). To further test whether reducing Dop2R signaling could rescue this severe phenotype, we introduced the *Dop2R* null allele [25] into the *PIGA* heterozygous null background (Fig 5B). In females lacking a single copy of *PIGA* and *Dop2R* (females have one mutant and one WT copy), recovery following seizure stimulation was dramatically improved. Female heterozygous *PIGA* null flies displayed a pronounced, long recovery deficit following mechanical stimulation, with only 15% leaving the bottom of the vial within 30 seconds while 52.9% of *PIGA* and *Dop2R* double mutant flies recovered in the same time frame (Fig 5B, S4 and S5 Videos). Males carrying the *Dop2R* hemizygous null allele and heterozygous *PIGA* null allele showed a similar improvement, increasing from 22.4% to 55% of flies climbing off the bottom at 30 seconds. We found a similar rescue effect with the *DAT* mutant (autosomal) [30]. Female *PIGA* heterozygous nulls improved from 20.3% to 91.7% off bottom at 30 seconds with *DAT* heterozygosity, and males improved from 31.7% to 83.1% (Fig 5C, S6 and S7 Videos). Together, these results demonstrate that modulating dopamine signaling, either by reducing receptor activity or limiting reuptake, can substantially rescue the severe seizure phenotype in the PIGA-CDG model, consistent with the improvements we observed in the *PIGA* eye model when dopamine signaling was similarly manipulated.

## Discussion

PIGA-CDG is a complex neurodevelopmental disorder with no disease-specific treatment. Because GPI-anchored proteins (GPI-APs) are broadly expressed and functionally diverse, it remains unknown how the loss of GPI-APs contributes to the neurological symptoms observed in patients. Although some symptoms of PIGA-CDG can be managed, the associated seizures are often severe and resistant to conventional treatments [3, 4, 35]. Identifying novel therapeutic compounds and molecular targets is crucial for enhancing seizure control and improving the overall quality of life for individuals with PIGA-CDG. Our lab leverages drug repurposing to identify small molecules that can quickly be translated from bench to bedside, circumventing many of the hurdles of *de novo* drug development. This strategy reveals candidate therapeutics for rare diseases, while shedding light on disease-modifying pathways and molecular mechanisms underlying the disorder.

In this study, we identified multiple FDA/EMA-approved drugs that improve a *Drosophila* model of PIGA-CDG. Utilizing what is known about the molecular targets of these compounds, we were able to use *Drosophila* genetic tools to validate modifier pathways underlying disease phenotypes in multiple models of PIGA-CDG. Of the twelve primary drug screen hits tested, we validated six drugs (50%) in dose-response experiments. Genetic validation of drug targets revealed cyclooxygenase (COX) inhibition and dopamine signaling as candidate therapeutic strategies for PIGA-CDG patients.

We show that three COX inhibitors tested in our *PIGA* eye model increased eye size, and that genetic knockdown of COX-like enzyme Pxt mimics this suppressor effect (Fig 2). Our findings suggest that inflammatory signaling plays a role in PIGA-CDG pathology. Loss of GPI-APs in the *Drosophila* eye is likely to induce cellular stress as misprocessed proteins that fail to undergo GPI anchoring are targeted for degradation [36]. Accumulation of unanchored or misfolded proteins in the ER can trigger the unfolded protein response, leading to ER stress, which is an established activator of inflammatory signaling pathways [37]. In this context, COX inhibition may mitigate downstream inflammatory signaling triggered by ER stress, thereby suppressing tissue damage in the *PIGA* eye model.

Notably, COX inhibition has also been shown to partially rescue another CDG model in our laboratory [12], suggesting that COX inhibition may be a shared therapeutic target across CDGs. Because CDGs share overlapping clinical features, it is plausible that some underlying disease mechanisms overlap and may be driven by inflammatory pathways. For example, aberrant glycosylation has been linked to neuroinflammatory responses in neurodegenerative diseases, suggesting that disruption of glycosylation machinery can similarly lead to neuroinflammation in CDGs [38].

Our primary screen also identified dopamine signaling as a potential modifier of *PIGA* disease phenotypes. In the PIGA-CDG eye model, disease severity was sensitive to both receptor-mediated signaling and dopamine availability in the presynaptic neuron and at the synapse. Loss of *Dop2R* improves the eye model, while loss of *Dop1R1* worsens it, suggesting that Dop2R signaling is pathogenic and Dop1R1 signaling is protective. This is consistent with prior work demonstrating the opposite effects of these receptors on cAMP signaling dynamics [24]. Notably, disrupting cAMP signaling is detrimental to the *PIGA* eye model, emphasizing the importance of cAMP homeostasis in the eye model. This could explain why *Dop2R* loss improves eye size and *Pde4* loss does not.

Further, our findings indicate that increasing dopamine availability through L-DOPA supplementation or DAT inhibition can partially rescue the PIGA eye model, supporting dopamine modulation as a candidate therapeutic option for PIGA-CDG patients. Interestingly, one PIGA-CDG patient has already been treated with Levodopa (L-DOPA), which alleviated his dyskinesia, improving quality of life [39].

The same genetic manipulations of the dopaminergic pathway that improved eye morphology also improved neurological phenotypes. Genetic loss of Dop2R improved the climbing deficit in the neuronal *PIGA* knockdown model and improved seizure recovery in the *PIGA* heterozygous null flies. Similarly, genetic loss of *DAT* also improved seizure recovery, reinforcing the idea that dopamine levels contribute to the severity of *PIGA* disease phenotypes. In contrast to what we observed in the eye, pharmacological inhibition of Dop2R using multiple D2R antagonists did not modify seizure phenotypes (data not shown). This could be because genetic inhibition of Dop2R is constant throughout development and could rewire neural circuits long term, while acute drug use is sensitive to off-target effects and inability to reach essential cell types that regulate behavioral changes. Drugs can degrade in food over time and likely have short half-lives compared to sustained, genetic loss of Dop2R, which may be what sets the *PIGA* neurological models apart from the *PIGA* eye model. Together, these suggest that the timing and specificity of Dop2R inhibition is important for attenuating *PIGA*-associated seizures. These findings indicate that the sensitivity to dopaminergic modulation in the *PIGA* eye model extends to neurological function, highlighting that dopaminergic dysregulation contributes to both developmental and neurological *PIGA* phenotypes.

With over 150 different GPI-APs, it is unclear how the loss of one or a few may impact dopamine synthesis and signaling. While components of the dopamine pathway are not GPI-anchored, multiple GPI-APs are known to play a role in neural circuitry, synaptic connectivity and plasticity, as well as axon guidance and regeneration [40]. Examples include Neural cell adhesion molecule (NCAM120), Glypicans, Contactins (CNTNs), and Neuroligins, all of which are critical to neurological function and are associated with various neurological disorders [41–43]. Supporting this, our lab identified a segregating loss of function variant in a PIGA-CDG pedigree with reduce penetrance [18]. Our lab further demonstrated that loss of CNTN2, a GPI-anchored cell adhesion molecule important for neural development, modifies developmental and neurological phenotypes in multiple *Drosophila* models of PIGA-CDG [18]. Disruption of neuronal GPI-APs may alter dopaminergic network organization and function, providing a potential mechanistic link between GPI-AP dysfunction and dopamine signaling. However, the contribution of any neural GPI-AP to dopamine signaling remains largely unexplored [40]. Loss of *Dop2R* improved phenotypes across three independent models, highlighting a potential mechanistic link between GPI-AP dysfunction and altered dopaminergic networks.

These findings parallel recent work from our lab describing how dopamine signaling and synthesis can modify DPAGT1-CDG, an N-linked glycosylation disorder. Similar to this study, Dop2R inhibition and increasing dopamine synthesis improved the DPAGT1-CDG model [12]. Perhaps this makes sense since RNAi knockdown of *PIGA* can rescue loss of *DPAGT1* in the *Drosophila* eye model and in cell culture, suggesting that overlapping disease pathogenesis might also indicate overlapping therapies [44]. This overlap reinforces the finding that disruptions in one glycosylation pathway can modify phenotypes arising from the perturbation of another glycosylation pathway [45].

One mechanistic possibility is that independent disruptions in glycosylation can converge on shared downstream signaling networks, such as dopamine signaling. For example, disruptions in GPI-anchor biosynthesis could reduce the surface expression of neuronal proteins that regulate dopamine signaling. In this context, loss of DPAGT1 could similarly disrupt proteins that regulate dopamine signaling, which can result in rebalancing of signaling and compensation when PIGA function is reduced. To date, our lab has demonstrated that dopaminergic signaling modifies disease phenotypes in three CDG models, including PIGA-CDG, DPAGT1-CDG, and NGLY1-deficiency [12, 13]. Together, these findings suggest that targeting dopaminergic signaling may represent a generalizable therapeutic strategy across CDGs.

Here, we screened 1,520 small molecules and identified several compounds that improved phenotypes in a *Drosophila* model of PIGA-CDG. Through genetic manipulation of the known therapeutic targets of these drugs, we uncovered new insights into PIGA-CDG pathophysiology, including links to COX-mediated inflammation and dopamine synthesis and signaling. Furthermore, we show that modulating dopamine signaling in patient-relevant PIGA-CDG models partially rescued disease-associated phenotypes, including seizures and locomotor deficits. Together, these findings not only highlight potential therapeutic pathways for PIGA-CDG but also suggest that more potent or selective D2R or COX inhibitors may provide enhanced efficacy in improving quality of life.

## METHODS

### Fly Stocks and Maintenance

Fly stocks were maintained at room temperature. Flies for model development and genetic validation work were obtained from the Bloomington Drosophila Stock Center (BDSC). All flies were fed with Glucose medium (D2) (Archon Scientific). F1 progeny generated by crossing *PIGA* RNAi (BDSC 62696) to *eya* composite-GAL4 were used for the primary drug repurposing screen and subsequent drug validation experiments, with all drug repurposing and validation experiments conducted in a 27°C incubator. We received *eya* composite-GAL4 as a gift from Justin Kumar (Indiana University Bloomington).

### Drug Repurposing Screen

We used the Prestwick Chemical Library (PCL), a collection of 1,520 FDA- or EMA-approved drugs. The PCL is divided into 19 plates, each containing 80 compounds. We are blinded to the identity of the compound during food preparation and analysis. We dilute compounds to 10 mM in DMSO, then further dilute 1:10 in 1X phosphate-buffered saline (PBS) for a final concentration of 1 mM. To prepare food, we use 500 cc bags of standard Glucose medium (D2) (Archon Scientific) and dispense it into a beaker. The food is microwaved until liquefied, then cooled on a heated stir plate until it reaches 60-70°C. While the food cools, we dispense 5 µL of each small molecule into test tube vials. We also prepare vials containing 5 µL of 1:10 DMSO:PBS solvent to serve as a control. For each plate of 80 drugs, we prepare 6-10 vials of solvent control vials. 1 mL of cooled liquid food is added to the aliquoted drug vials and solvent controls for a final concentration of 5 µM drug and 0.05% DMSO, then vortexed to mix. Food containing a drug is allowed to cool for at least 2 hours.

Once cooled, 4-6 premated *PIGA* RNAi females and 3-4 *eya* composite-GAL4 males are added to each vial. Flies were allowed to lay eggs for 1-4 days at 27°C. Parent flies were discarded when vials contained 45-60 eggs. Both females and males are homozygous, so all F1 progeny carried the *eya* composite-GAL4 and *PIGA* RNAi. All flies were collected from the drug vials at 2-4 days post-eclosion and frozen at -20°C. Up to 5 males (or females if males were not collected) per drug were imaged at 3x magnification (Leica EC3 camera). To quantify eye size, eyes were manually traced using a tablet, and the area was measured (ImageJ). To maintain consistency, only the left eyes were imaged and quantified. Z-Scores were calculated for each drug treatment by subtracting the average eye area of flies on drug from the average eye area of flies on DMSO across all controls, divided by the standard deviation of eye area across all controls (S1 Table). Z-Scores were then rank-ordered, and we defined compounds that suppressed the eye phenotype as Z ≥ 1.5 and compounds that enhanced the eye phenotype as Z ≤ -1.5 (S2 Table) [12, 19].

### Drug Validation

To account for potential differences in drug formulation, we obtained compounds from secondary sources, including MedChemExpress and Cayman Chemical (S2 Table). Instead of resuspending validation drugs in 1:10 DMSO:PBS, we dissolved the drugs in solvents that allowed us to achieve high enough solubility to test at final concentrations of 1 µM, 5 µM, 25 µM, and in some cases, 50 µM. Due to potential technical variations in food preparation and drug formulation, we consider a drug to be validated when eye size is increased at any of these doses when compared to the *PIGA* eye model raised on the respective solvent control. Compounds and their respective vehicles are listed in S2 Table. 6-31 male flies were imaged and quantified for each drug or solvent condition.

Percentages listed in the text and S2 Table represent the percentage change of the treated mean compared to the control mean. Raw pixel values of flies treated with drug or control solvent can be found in S4 Table.

### Genetic Validation

To introduce additional RNAi constructs or mutant alleles in the *PIGA* model, we used w-; *PIGA* RNAi/CyO; *eya*-GAL4 flies. All genetic validation crosses for the *PIGA* eye model were done in a 20°C incubator. 7-10 male flies were imaged and quantified for each genetic condition.

To modify dopamine signaling synthesis, we used virgin females harboring a *Dop2R* null allele (BDSC 84720), *Dop1R1* null allele (BDSC 92640), *Dop1R2* null allele (BDSC 84719), *ple^4^* null allele (DGGR 106-955), *Ddc* null allele (BDSC 3190), *Vmat* null allele (BDSC 29477), *DAT^Z2−1744^* null allele (BDSC 30867), *Pde4^dnc^* null allele (BDSC 6020), or *Adcy1^rut-1^* null allele (BDSC 9404) and crossed them to w-; *PIGA* RNAi/CyO; *eya*-GAL4 males (and *eya* composite-GAL4 controls), selecting for non-CyO flies.

To investigate how loss of prostaglandin synthesis impacted the *PIGA* eye model, we used virgin females of *Pxt* RNAi (BDSC 28934) and *Pxt* RNAi (BDSC 32382) and crossed them to w-; *PIGA* RNAi/CyO; *eya*-GAL4 males (and *eya* composite-GAL4 controls).

RNAi crosses used either attP40 (BDSC 36304) or attP2 (BDSC 36303) as control comparisons, and null crosses used w^1118^ (VDRC 60000) or Canton-S. Canton-S was a gift from Carl Thummel.

Percentages listed in the text and S3 Table represent the percentage change of the treated mean compared to the control mean. Raw pixel values of the control *PIGA* eye model or genetically modified *PIGA* eye model can be found in S4 Table.

### Negative Geotaxis Assay

Flies were raised at 25°C. 10-15 female flies (3-5 days old) were separated into a vial, off CO2 for over 24 hours. Flies were tapped to the bottom of the vial and allowed to climb. At 30 seconds, the proportion of flies climbing was scored by the number of flies on the bottom or off the bottom of the vial. All climbing experiments were performed with at least 55 flies per genotype. For negative geotaxis assays, virgin females (BDSC 84720 or BDSC 30867) were crossed to w-; *PIGA* RNAi/CyO, GAL80; *elav*-GAL4 males, selecting against CyO. Neuronal PIGA knockdown flies were generated by crossing *PIGA* RNAi (BDSC 62696) virgin females to *elav*-GAL4 (BDSC 46655) males. Heterozygous mutant flies were generated by crossing virgin females (BDSC 84720 or BDSC 30867) to *elav*-GAL4 males. Only female flies were used for the Negative Geotaxis Assay, as male flies exhibited severe impairment that prohibited reliable assessment of climbing performance.

### Bang Sensitivity Assay

Flies were raised at 25°C. 10-15 flies (3-5 days old) were separated into a vial, off CO_2_ for over 24 hours. Flies were transferred to experimental vials without food and immediately vortexed at maximum intensity for 10 seconds. The proportion of flies recovered at 30 seconds was calculated as the number of flies that left the bottom of the vial divided by the total number of flies tested. For bang sensitivity assays, virgin females (BDSC 84720, BDSC 30867, or w^1118^ control) were crossed to *PIGA* heterozygous null flies previously characterized, selecting against CyO, GFP [16]. Male and female flies were tested separately.

### Statistics

For all drug and genetic experiments on the *PIGA* eye model, we calculated statistical significance using One-way ANOVA followed by Dunnett’s test to compare between individual groups and correct for multiple comparisons. All preprocessing was performed in R Studio using the dplyr package. Due to technical variability associated with drug formulation, dosing, and inconsistent ingestion, we identified outliers within each drug treatment group using the IQR method (1.5×IQR rule) and excluded them prior to statistical analysis. In contrast, genetic phenotypes reflect stable, developmentally integrated perturbations, so variation in those datasets is considered biologically meaningful and retained. For all behavioral experiments, we calculated statistical significance using a Chi-squared test. We used R for these analyses.

## ACKNOWLEDGMENTS

We dedicate this work to Emmett Nguyen, the first boy we met with PIGA-CDG. His life and parents, Ann and Steve, inspired us to begin work on this rare disorder. We thank Dr. Katherine Beebe, Emily Coelho, Delaney Baratka, and Caroline Massey for their technical advice regarding the drug repurposing screen. We thank Elsie Smith for her assistance in the lab. We thank Dr. Hugo Bellen and Dr. Oguz Kanka for generating and sharing the *PIGA* null allele. A special thank you to Dr. Holly Thorpe for advising on all things *PIGA* and her unwavering support throughout the course of this work and manuscript preparation.

