## Supplemental Figure 1 for "A drug repurposing screen reveals dopamine signaling as a candidate therapeutic pathway for PIGA-CDG"

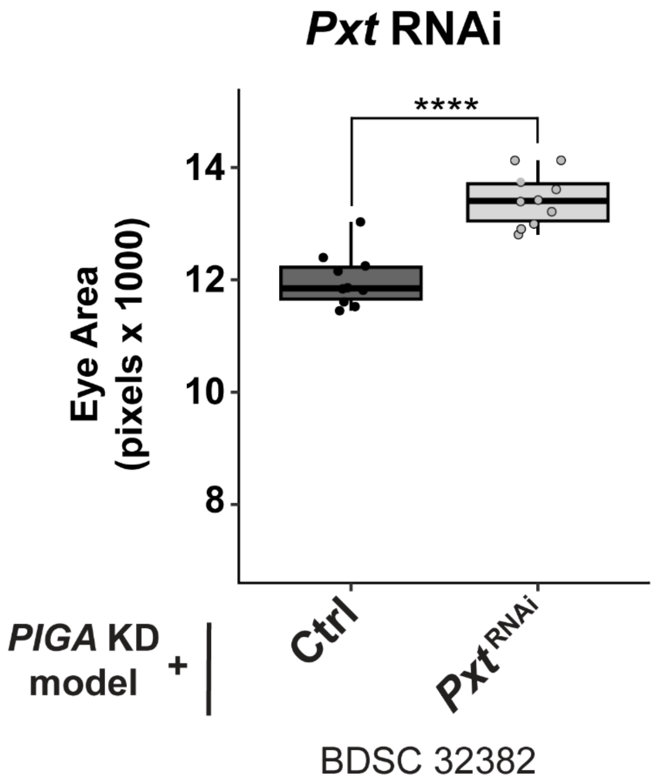


**S1 Fig: Secondary RNAi validation for Pxt.** RNAi against COX-like gene *Pxt* (BDSC 32382) partially rescues eye size of *PIGA* eye model. Statistical significance was determined using unpaired two-tailed t-test. **** p<0.0001.
