## Supplemental Figure 2 for "A drug repurposing screen reveals dopamine signaling as a candidate therapeutic pathway for PIGA-CDG"

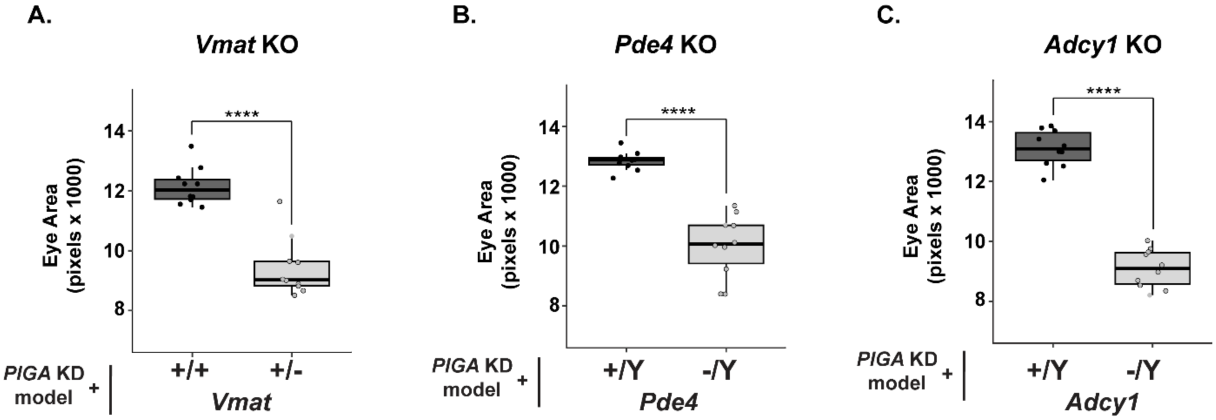


**S2 Fig: Loss of monoaminergic trafficking and dysregulation of cAMP signaling is detrimental to *PIGA* eye model. (A)** Introducing a heterozygous null allele of *Vmat* (BDSC 29477), which encodes for the enzyme responsible for transporting monoamine neurotransmitters into synaptic vesicles, reduces *PIGA* eye size by 22.4%. **(B)** A hemizygous null allele of *Pde4* (BDSC 6020), a gene that encodes a cAMP phosphodiesterase, decreases eye size by 22%. **(C)** A hemizygous null allele of *Adcy1* (BDSC 9404), a gene that encodes an adenylate cyclase which synthesizes cAMP, decreases eye size by 30.6%. Statistical significance was determined using unpaired two-tailed t-tests for genetic experiments. * $p<0.05$, ** $p<0.01$, *** $p<0.001$, **** $p<0.0001$.
