## Supplemental Figure 3 for "A drug repurposing screen reveals dopamine signaling as a candidate therapeutic pathway for PIGA-CDG"

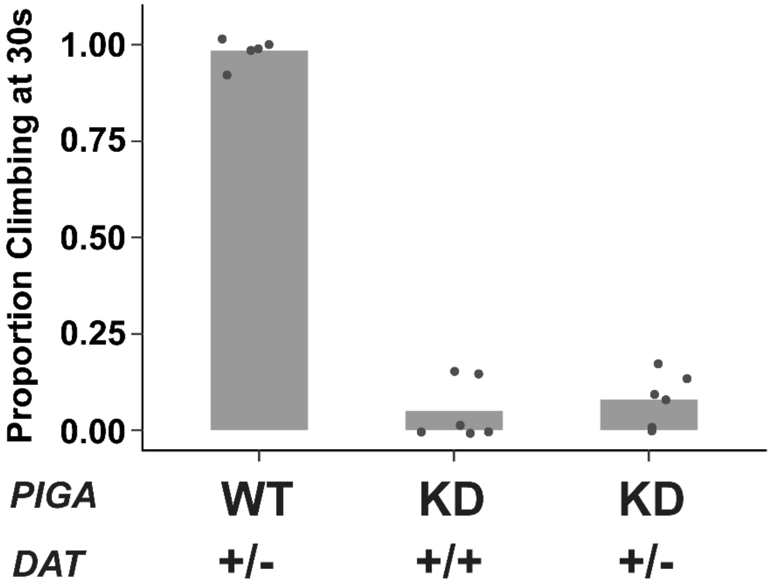


**S3 Fig. *DAT* loss does not impact pan-neuronal *PIGA* knockdown climbing phenotype.** Heterozygous loss of *DAT* improves the severe climbing impairment in pan-neuronal *PIGA* knockdown female flies. Only females were tested, as males were too severely impaired to be properly assessed.
